# Loss of the extracellular matrix protein DIG-1 causes glial fragmentation, dendrite breakage, and dendrite extension defects

**DOI:** 10.1101/2021.08.12.456129

**Authors:** Megan K. Chong, Elizabeth R. Cebul, Karolina Mizeracka, Maxwell G. Heiman

## Abstract

The extracellular matrix (ECM) guides and constrains the shape of the nervous system. In *C. elegans*, DIG-1 is a giant ECM component that is required for fasciculation of sensory dendrites during development and for maintenance of axon positions throughout life. We identified four novel alleles of *dig-1* in three independent screens for mutants affecting disparate aspects of neuronal and glial morphogenesis. First, we find that disruption of DIG-1 causes fragmentation of the amphid sheath glial cell in larvae and young adults. Second, it causes severing of the BAG sensory dendrite from its terminus at the nose tip, apparently due to breakage of the dendrite as animals reach adulthood. Third, it causes embryonic defects in dendrite fasciculation in inner labial (IL2) sensory neurons, as previously reported, as well as rare defects in IL2 dendrite extension that are enhanced by loss of the apical ECM component DYF-7, suggesting that apical and basolateral ECM contribute separately to dendrite extension. Our results highlight novel roles for DIG-1 in maintaining the cellular integrity of neurons and glia, possibly by creating a barrier between structures in the nervous system.

## INTRODUCTION

Neurons and glia exhibit a kaleidoscopic array of shapes, with each cell type adopting a distinctive morphology that is intimately coordinated with its immediate neighbors [1]. Most studies of cell morphogenesis have focused on factors that sculpt cells from the inside, especially regulators of the cytoskeleton. By contrast, unbiased genetic approaches have pointed to an equally important role for factors that act from outside the cell – in particular, components of the extracellular matrix (ECM).

*C. elegans* provides a powerful system to identify novel factors that shape neurons and glia. Briefly, the morphology of every neuron and glial cell is highly stereotyped and has been carefully catalogued; individual cells can be readily visualized in live intact animals throughout life; and it is straightforward to perform forward genetic screens by visually identifying mutants that show aberrant morphology of single neurons or glial cells. Recent screens have revealed roles for extracellular factors in diverse aspects of neurodevelopment, including neuroblast migration [2–4], axon and dendrite extension [5–9], fasciculation of neurites into nerve bundles [10–15], and synapse development [16–21].

By examining *C. elegans* sense organs, we previously identified several extracellular factors that control neuronal and glial morphogenesis [9,10,22–26]. We initially focused on the major sense organ, the amphid, which consists of 12 sensory neurons and two glial cells, called the amphid sheath – which is examined further in this study – and the amphid socket [27]. The neurons extend unbranched sensory dendrites to the nose tip, where most of them protrude into the external environment through a pore formed by the sheath and socket glia [27]. We showed that this structure develops embryonically as a narrow epithelial tube lined by the apical ECM protein DYF-7 [9]. In the absence of DYF-7, this tube ruptures, causing the sensory neurons and sheath glial cell to detach from the nose and resulting in severely shortened dendrites [9,22].

In other sense organs, sensory dendrites protrude through pores in their own dedicated glia. For example, the IL2 neurons – another cell type examined in this study – protrude through pores in IL sheath and socket glia. We previously found that these neurons also require DYF-7 for dendrite extension, suggesting that they similarly develop as part of an epithelial tube [9]. By contrast, some sensory dendrites extend to the nose tip but do not pass through a glial pore and do not require DYF-7. For example, the carbon dioxide-sensing neuron BAG – the third cell type examined in this study – forms elaborate membranous attachments along the surface of a specific glial partner but does not enter a glial pore [9]. Unlike the amphid and IL2 neurons, BAG does not require DYF-7 for dendrite morphogenesis, but instead requires a cell adhesion molecule that acts both in the neuron itself and in glia [23]. Together, these examples illustrate how neurons and glia can be shaped by extracellular factors that mediate specific physical interactions.

Here, we report that three genetic screens for mutants affecting disparate cellular phenotypes – amphid sheath glial cell morphology, IL2 dendrite extension, and BAG dendrite integrity –all identified lesions in the same gene, encoding the giant ECM molecule DIG-1. DIG-1 was previously shown to affect neuronal cell body positioning and the fasciculation of axons and dendrites, including amphid and IL2 dendrites [11,12,28]. We find that loss of DIG-1 also causes a novel glial fragmentation defect, breakage of BAG dendrites, and detachment of IL2 dendrites from the nose tip. Glial fragmentation and BAG dendrite breakage occur in late larval and early adult stages, while IL2 dendrite defects appear to arise during embryonic development and are enhanced by loss of DYF-7, which also acts embryonically. These observations raise the question of how a single ECM component affects such diverse neuronal and glial cell types with distinctive outcomes and different developmental timing. We consider the possibility that DIG-1 protects the structural integrity of neurons and glia by creating a barrier between them and their surroundings.

## RESULTS

### Genetic screens for disparate glial and neuronal phenotypes identify lesions in DIG-1

To identify factors that control the morphology of glia and neurons, we performed visual forward genetic screens using cell-type-specific fluorescent markers. Briefly, we mutagenized strains bearing markers for the amphid sheath glial cell (*F16F9.3*pro:mCherry), the sensory neuron BAG (*flp-17*pro:GFP), or the sensory neuron IL2 (*klp-6*pro:GFP); allowed animals to self-fertilize for two generations to yield F2 progeny bearing random homozygous mutations; and visually screened non-clonal pools of progeny using a fluorescence stereomicroscope to isolate individuals with aberrant cell morphology (Fig. 1A). Four mutants (*hmn152, hmn158, hmn227,* and *hmn259*) in which the phenotype of interest was transmitted with high penetrance across generations were analyzed further (Fig. 1A). All four mutants were recessive, and we used whole-genome sequencing of pooled recombinants to identify possible causative mutations.

**Figure 1.**
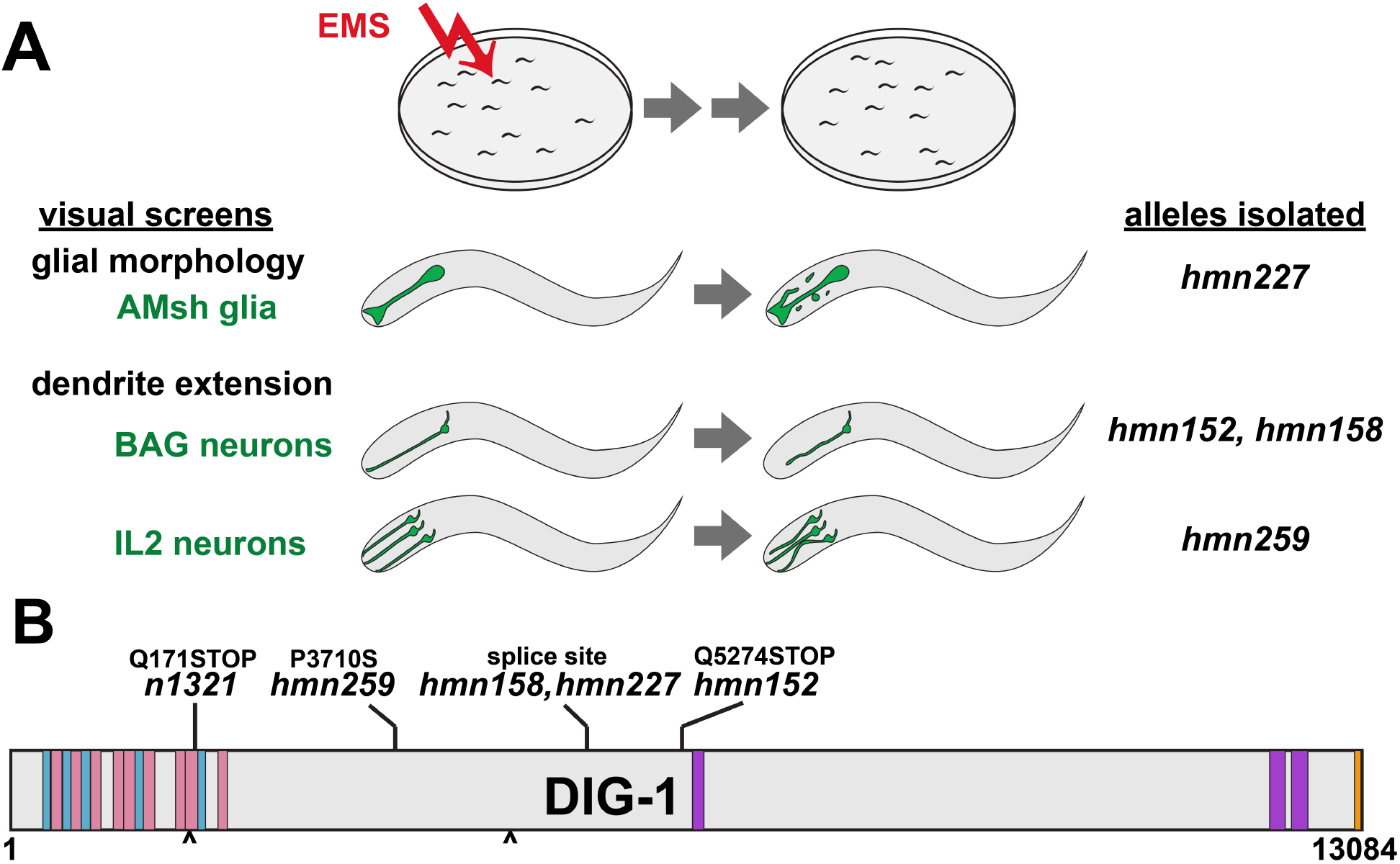
Overview of genetic screens and *dig-1* alleles. (A) Schematic of screening strategy, phenotypes, and resulting alleles isolated. Animals bearing markers for the amphid sheath (AMsh) glia, BAG neurons, or IL2 neurons were mutagenized with ethyl methanesulfonate (EMS), grown for two generations, and screened for morphology defects, yielding four alleles of *dig-1*. (B) Schematic of DIG-1 protein depicting the nature of each newly isolated allele as well as the reference allele *n1321* used throughout this study. Blue, Ig domains; pink, FN domains; purple, VWA domains; orange, EGF domain; carets, RGD motifs.

Lesions in *dig-1* were identified in mutants from each screen (Fig. 1B). *dig-1* is predicted to encode a 13,084 amino acid ECM protein consisting of an amino-terminal region comprised of repetitive Immunoglobulin (Ig)-like and Fibronectin III (FN)-like domains that suggest roles in cell adhesion, a large central region of poorly-conserved sequence with unknown function, and a carboxy-terminal region containing von Willibrand Factor A (VWA) and epidermal growth factor (EGF)-like domains that are often found in ECM and cell adhesion proteins (Fig. 1B) [11,12]. It contains two RGD motifs, suggesting a possible role in integrin binding (Fig. 1B). Three of the alleles we isolated are predicted to disrupt a splice site (*hmn158* and *hmn227;* these were isolated in independent screens but result in an identical nucleotide change), or introduce a premature termination codon (*hmn152*), suggesting they may represent severe loss of function mutations (Fig. 1B). Interestingly, one allele (*hmn259*) introduces a Pro>Ser missense mutation at position 3710 in a poorly conserved region, pointing to a possible functional role for this otherwise unannotated sequence (Fig. 1B). Interestingly, a previously reported allele, *nu336*, introduces a Ser>Phe missense mutation in approximately the same region, at position 3988 [11]. It is worth noting that both of these alleles were isolated in screens for IL2 morphogenesis defects.

Three lines of evidence argue that *dig-1* lesions are causal for the phenotypes we identified. First, sequencing of pooled recombinants showed that the lesions are genetically linked to the phenotypes. Second, the phenotypes were recapitulated using the reference allele *dig-1(n1321),* which is predicted to introduce a premature termination codon at position 171 [12,29]. Third, the reference allele failed to complement each of the alleles for their respective phenotypes (*n1321/hmn227*, amphid sheath defects in 10/22 fourth larval stage (L4) animals; *n1321/hmn152* and *n1321/hmn158*, BAG defects in 5/27 and 9/24 L4 animals, respectively; *n1321/hmn259,* IL2 defects in 8/17 L4 animals). We did not perform transgenic rescue experiments due to the size of the *dig-1* gene and cDNA (48 kb and 39 kb respectively). However, taken together, the linkage, recapitulation, and non-complementation results strongly suggest that the *dig-1* lesions are causative for these phenotypes. For consistency, we used the reference allele *dig-1(n1321)* for all further experiments in this study.

### Loss of DIG-1 causes glial fragmentation in young adults

The amphid is the largest sense organ in *C. elegans*, consisting of 12 neurons and the sheath and socket glia [27]. The sheath develops in concert with the neurons, together forming a multicellular rosette in the embryo that re-organizes during embryo morphogenesis to form an epithelial tube in which the sheath glial cell wraps the dendrite endings [9,25]. In the mature structure, the amphid sheath cell body is located near those of the neurons and extends a process collateral with the amphid dendrites, terminating at the nose where it adopts a characteristic wing-shaped morphology [27]. The dendrites of the 12 amphid neurons penetrate into the sheath, with the sheath forming ring-shaped tight junctions around each dendrite individually [9,27,30]. Four neurons (AWA, AWB, AWC, AFD) form elaborate ciliated endings that are embedded in the wing-shaped portion of the sheath, while the others (ASE, ADF, ASG, ASH, ASI, ASJ, ASK, ADL) terminate in simple cilia that lie in a central lumenal channel [27,30]. Previous genetic screens have identified mutants that affect amphid sheath specification and cell body positioning [31], that disrupt extension of the amphid sheath glial process [9,22], or that lead to enlarged or supernumerary amphid sheath cells [32,33].

In *dig-1* mutant animals, we observed a defect that, to our knowledge, has not been seen in other mutants affecting amphid sheath development. While the amphid sheath appears superficially normal in newly hatched first larval stage (L1) animals, it exhibits progressively more severe disorganization in L4 and two-day adult (2A) animals (Fig. 2). The characteristic wing-shaped ending becomes enlarged, extending more posteriorly along the head (Fig. 2B,C). Abnormal protrusions or empty spaces (vacuoles) are often seen at the cell ending (Fig. 2B,C). Most strikingly, the glial cell appears to fragment, with a multitude of ~1-5 μm fluorescent blebs appearing around the sheath glial process (Fig. 2B,C). The distribution of these fragments suggests they may be internalized by neighboring cells. Interestingly, the amphid sheath was recently shown to internalize extracellular vesicles released by amphid neurons and, in the absence of the sheath, other neighboring cells perform this function, suggesting that several cells in the head are able to internalize cellular debris [34]. This phenotype is highly penetrant, with >80% of individual amphid sheath glial cells showing extensive fragmentation (Fig. 2D).

**Figure 2.**
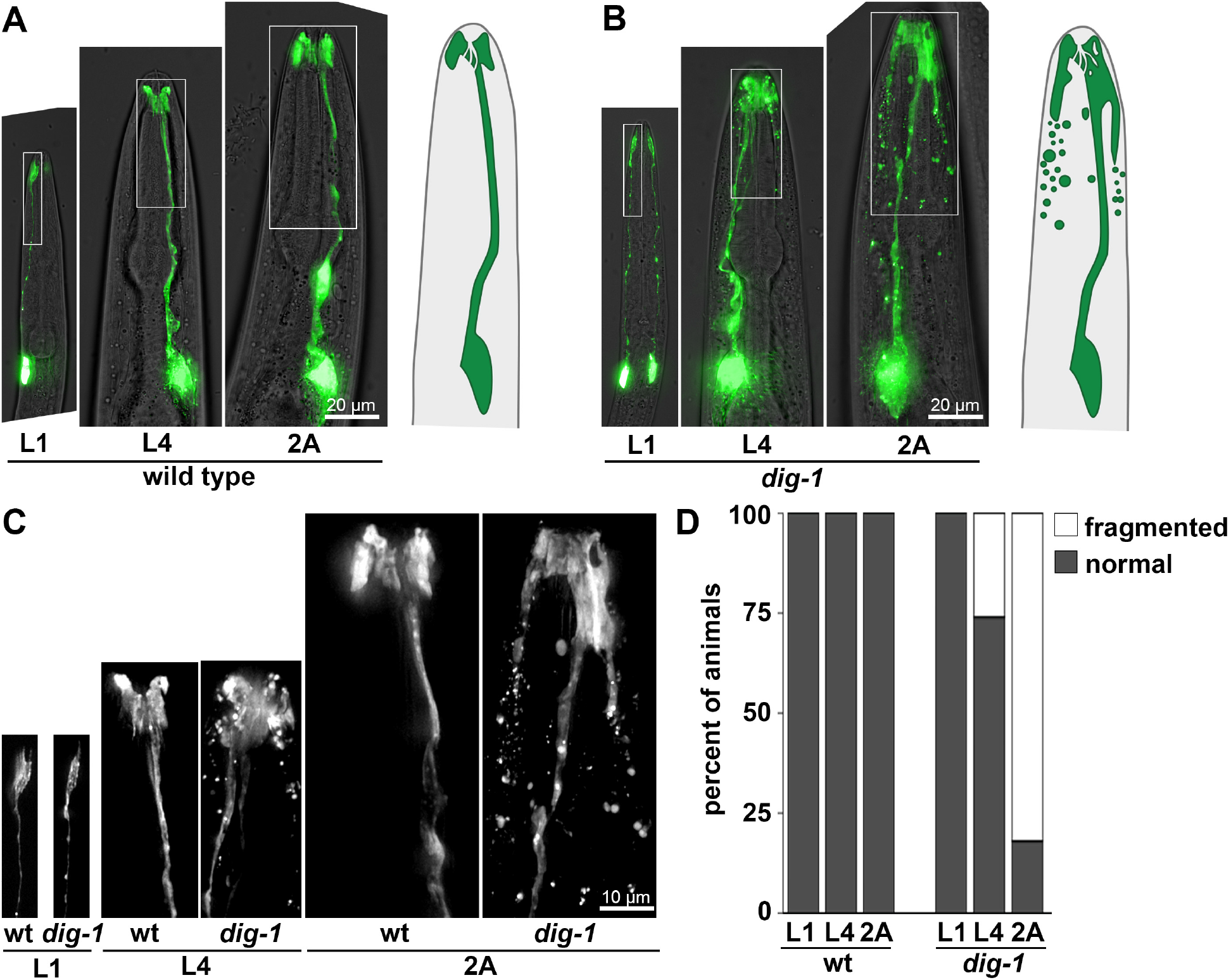
Amphid sheath glia undergo fragmentation in *dig-1* mutants. Heads of (A) wild-type and (B) *dig-1(n1321)* animals at the L1, L4, and 2A stages, showing progressive fragmentation of amphid sheath in L4 and 2A *dig-1* animals. Animals express *F16F9.3*pro:mCherry, pseudocolored green. Schematics of amphid sheath glia are shown on the right. (C) 2x magnification of boxed regions in A and B. (D) Quantification of phenotype penetrance at each stage.

### Loss of DIG-1 causes breakage of BAG dendrites in young adults

BAG is a carbon dioxide sensing neuron that forms a specialized attachment to a specific glial partner, the inner labial socket (ILso) glial cell [23,27]. Like other head sensory neurons, BAG extends an unbranched dendrite to the nose tip where it terminates in a sensory cilium. However, the BAG dendrite does not penetrate into a sheath glial cell. Instead, its cilium forms a membranous “bag” that precisely wraps a thumb-like protrusion of the ILso glial cell [23,27]. During development, the BAG dendrite extends by attaching near the presumptive nose tip and then stretching out during embryo elongation [23]. BAG dendrite extension requires the adhesion molecule SAX-7, which acts both in BAG and in glia, as well as the cytoskeletal organizer GRDN-1, which acts in glia [23]. In the absence of SAX-7 or GRDN-1, the BAG dendrite detaches from the nose during embryo elongation, resulting in severely shortened dendrites in the mature structure [23].

While screening for additional mutants that affect BAG dendrite extension, we isolated two alleles of *dig-1* that exhibited an unusual phenotype. Unlike *sax-7* and *grdn-1* mutants, in which BAG dendrites appear shortened from L1 through adulthood, *dig-1* mutants exhibited superficially wild-type BAG dendrites at the L1 stage. However, at later stages, these dendrites appeared to undergo constriction at a stereotyped region of the distal dendrite, and ultimately appeared to break, especially as animals reached adulthood (Fig. 3). In most cases, a brightly fluorescent remnant was still visible at the nose tip, suggesting that the specialized attachment of BAG to the ILso glial cell remained intact (Fig. 3B,C). The proximal dendrites often appeared wavy or curved, like a tightly-stretched string that has been cut (Fig. 3B,C). Overall, this phenotype was highly penetrant, with 72% of BAG dendrites appearing fully severed in 2A animals (Fig. 3D). An additional 18% of BAG dendrites had a thin, dimly fluorescent connection – often visible only at longer exposure settings – between the main dendrite and the remnant at the nose (Fig. 3B-D, “constricted”). We noted that wild-type BAG neurons appear to exhibit dimmer fluorescence in this region, suggesting they may become thinner here, possibly due to compressive force from surrounding structures (Fig. 3A,C). This raises the possibility that DIG-1 protects BAG dendrites from breakage at a naturally occurring pinch point.

**Figure 3.**
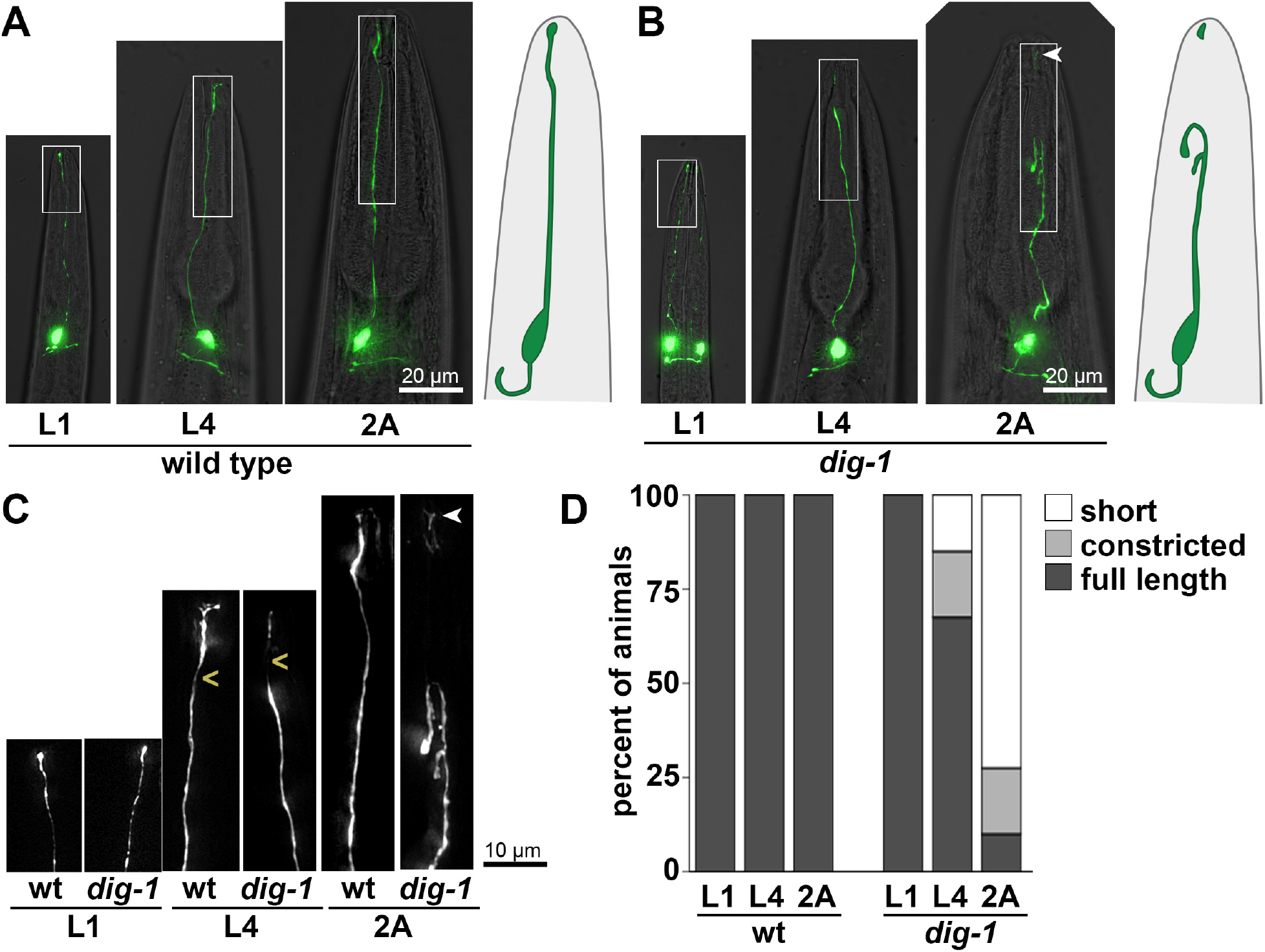
BAG dendrites constrict and break in *dig-1* mutants. Heads of (A) wild-type and (B) *dig-1(n1321)* animals at the L1, L4, and 2A stages, showing apparent breakage of the BAG dendrite in L4 and 2A *dig-1* animals. Animals express *flp-17*pro:GFP. Schematics of BAG neurons are shown on the right. (C) 2x magnification of boxed regions in A and B. (D) Quantification of phenotype penetrance at each stage. “Constricted” refers to dendrites in which a thin, barely visible connection persists between the main dendrite and the remnant at the nose tip, as shown in the L4 *dig-1* animal. This region (marked by <) appears thinner in wild-type animals as well. Arrowhead, remnant at the nose tip following dendrite breakage.

### Loss of DIG-1 causes defasciculation and dendrite extension defects in IL2 neurons

IL2 neurons are a set of six chemosensory neurons, arranged as dorsal, lateral, and ventral pairs [27]. Each IL2 dendrite penetrates into its sheath glial cell as part of an epithelial tube, with its ciliated ending exposed directly to the outside environment. Similar to amphid neurons, IL2 neurons show a complete dependence on the apical ECM molecule DYF-7 for dendrite extension and, in *dyf-7* mutants, 100% of IL2 neurons fail to reach the nose tip [9]. However, unlike amphid neurons, which become severely shortened in *dyf-7* mutants (~5-10% of wild-type length), IL2 neurons are more mildly affected (~80-90% of wild-type length) [9]. Therefore, to identify other factors that contribute to IL2 dendrite extension, we performed a visual screen for mutants with abnormal IL2 dendrites and identified *dig-1* as a mutant with two abnormal aspects of IL2 dendrite morphology.

First, as has been described previously, loss of DIG-1 causes pronounced defasciculation defects in IL2 dendrites (Fig. 4) [11]. Rather than extending directly to the nose tip in distinct parallel fascicles as in wild-type animals, IL2 dendrites took a meandering course in *dig-1* mutants, often appearing to fasciculate together for some distance before separating again (Fig. 4B,C). Other defasciculation defects have also been observed in *dig-1* mutants, including in amphid dendrites [10,11] and several axon tracts [12], suggesting DIG-1 plays a general role in maintaining neurite bundles. However, while other defasciculation defects appear progressively throughout larval growth, consistent with previous work we find that IL2 defasciculation defects are present from the L1 stage with little change in phenotype penetrance as animals grow (Fig. 4D) [11]. In addition to dendrite defects, IL2 neurons also exhibited variable cell body positioning at all stages, as reported previously [11]. It remains unclear why DIG-1 is required embryonically for some nerve bundles, and only post-embryonically for others.

**Figure 4.**
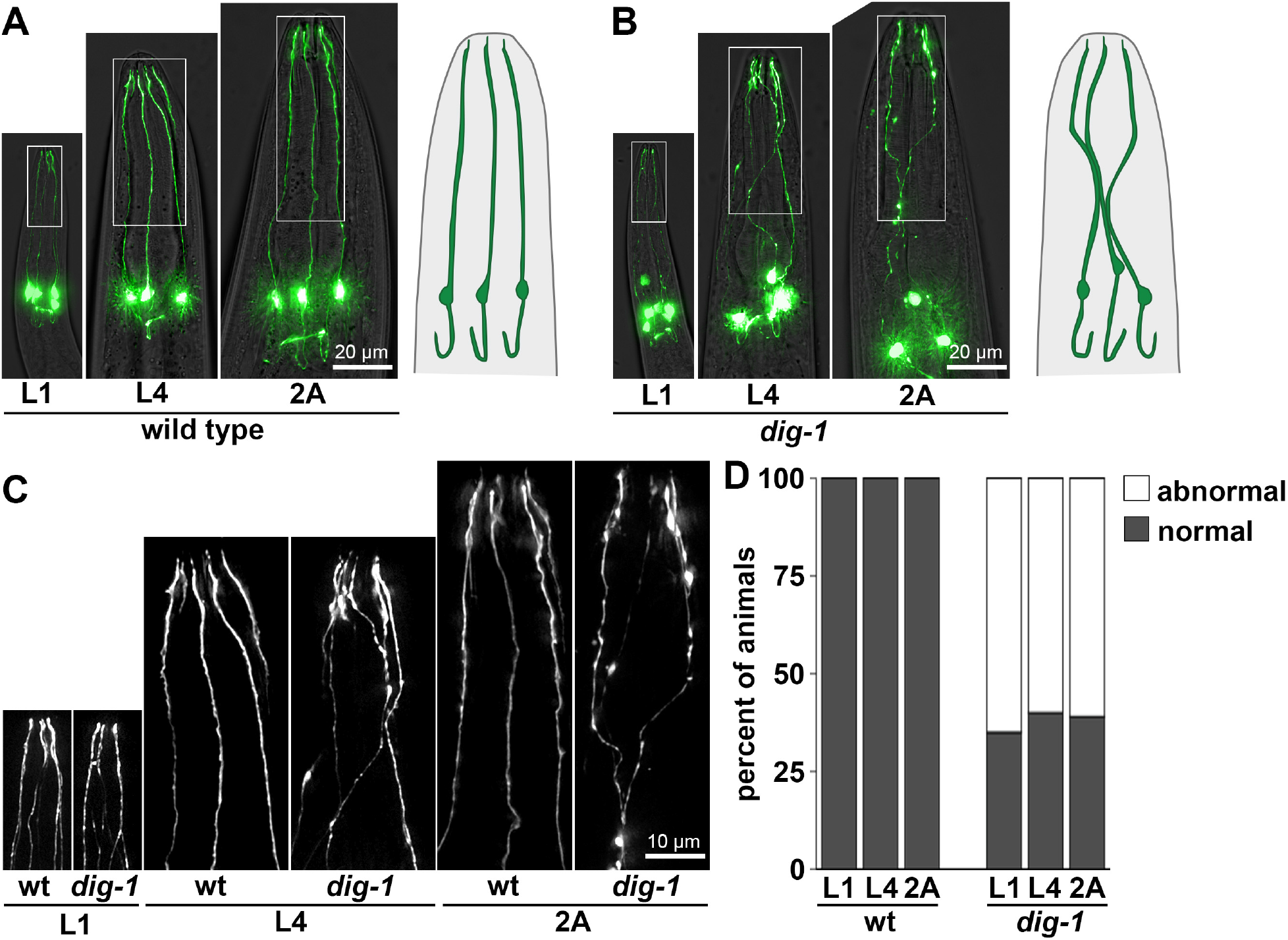
IL2 dendrites exhibit fasciculation defects in *dig-1* mutants. Heads of (A) wild-type and (B) *dig-1(n1321)* animals at the L1, L4, and 2A stages, showing fasciculation defects at all stages in *dig-1* animals. Animals express *klp-6*pro:GFP. Schematic of IL2 neurons are shown on the right. (C) 2x magnification of boxed regions in A and B. (D) Quantification of phenotype penetrance at each stage.

Second, we observed rare (6%) dendrites that were less than half their normal length (Fig. 5C,E), indicating a role for DIG-1 in dendrite extension. To test for a genetic interaction with *dyf-7,* we examined *dig-1*; *dyf-7* double mutants using the presumptive null allele *dyf-7(ns119)* which introduces a premature termination codon at position 3 [22]. We focused on L4 animals, because in L1 animals the smaller size and decreased brightness of dendrites combined with the fasciculation defects made it difficult to measure dendrite lengths accurately. In the *dig-1; dyf-7* animals, the average IL2 dendrite length was shorter and dendrite lengths were overall more variable than in either single mutant (Fig. 5D,E), suggesting that DIG-1 acts independently of DYF-7 – that is, there is a role for DIG-1 in promoting IL2 dendrite extension even in the absence of DYF-7 (Fig. 5). It is worth noting that DYF-7 is a component of the apical ECM, while DIG-1 is thought to primarily be present in basal ECM, suggesting that apical and basal ECM may contribute independently to sensory dendrite extension.

**Figure 5.**
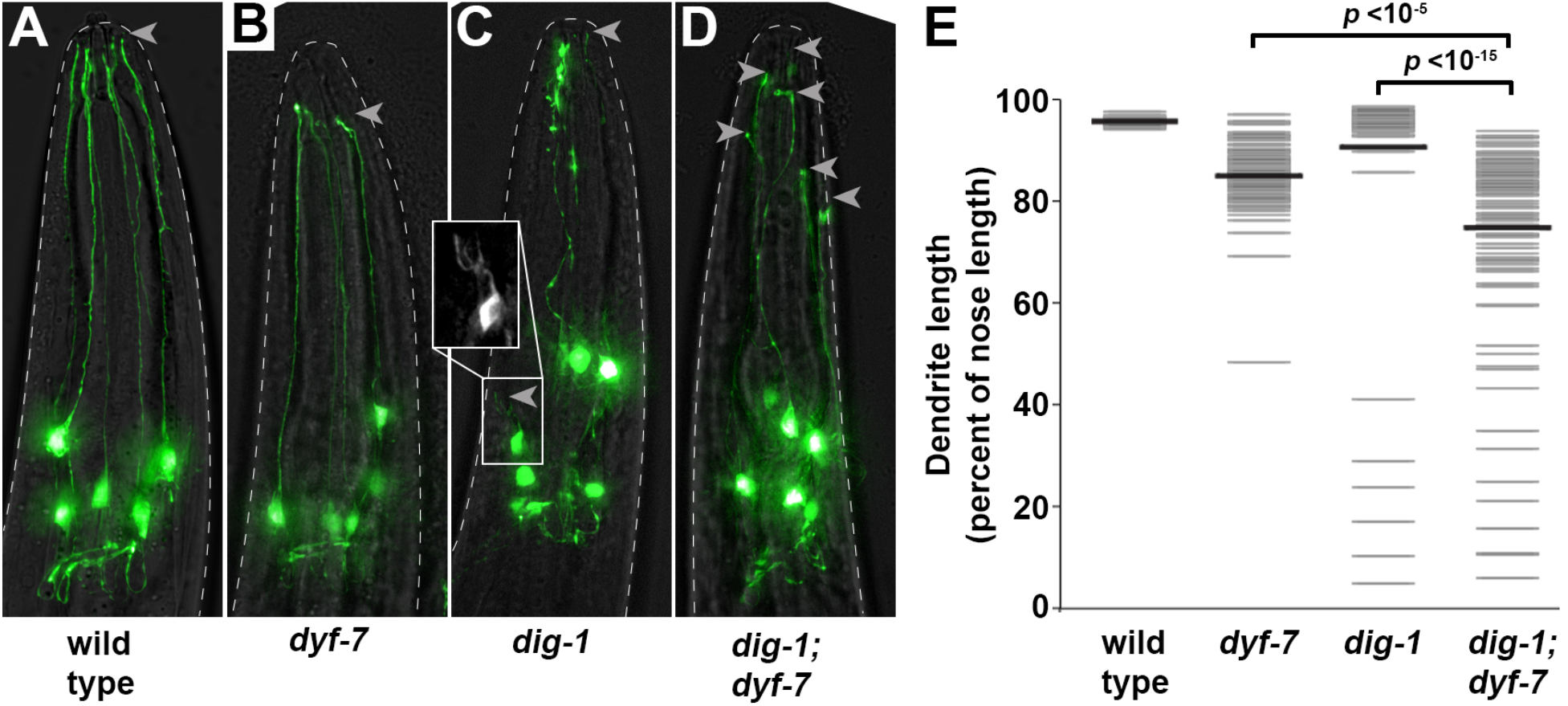
Rare IL2 dendrite extension defects in *dig-1* mutants are enhanced by loss of *dyf-7*. Heads of (A) wild-type, (B) *dyf-7(ns119)*, (C) *dig-1(n1321)*, and (D) *dig-1(n1321); dyf-7(ns119)* animals at the L4 stage, showing dendrite extension defects. Arrowheads, dendrite endings. A severely shortened dendrite is boxed and magnified in C. Animals express *klp-6*pro:GFP. (D) Quantification of phenotype expressivity. Dendrite lengths were measured as a fraction of the distance from the cell body to the nose tip. Gray bars, individual dendrites; black bars, population mean. *n*>90 per genotype. *p-*values, Mann-Whitney U test.

## DISCUSSION

The functions of ECM are only beginning to be understood. Here, we describe additional phenotypes resulting from loss of DIG-1, a giant ECM protein, that shed light on its roles in maintaining the integrity of the nervous system. Although the developmental mechanisms that produce these phenotypes remain unclear, each of them can be interpreted in terms of cellular breakage: fragmentation of the amphid sheath; constriction and breakage of the BAG dendrite near its ending; and detachment of IL2 dendrites in a manner similar to the rupture of glial epithelial tubes caused by loss of DYF-7 [9].

Previous phenotypes associated with DIG-1 had pointed most strongly to roles in adhesion. These include the displaced gonad defects for which *dig-1* is named [29]; defasciculation of amphid dendrite bundles and wandering of other dendrites including those of the IL2 neurons [10,11]; defects in neuronal cell body positioning [12,28]; and “flip-over” defects of several axons (for example: PVQ, AVK, and RMEV) in which axons are displaced across the midline and then return to their correct path [12]. Several of these phenotypes, including axon flip-over, arise progressively throughout larval development and are suppressed by inhibiting locomotion, suggesting they are caused by mechanical forces due to growth and movement [12]. One interpretation of these defects is that DIG-1 normally acts as an adhesion protein to keep neurites tightly attached to their correct partners.

Our results suggest a different, but not mutually exclusive, role in which DIG-1 creates a barrier between tightly packed structures in the nervous system. For example, DIG-1 may contribute structurally to the overall basal ECM that separates cells. Alternatively, it may form a coat around the extracellular surfaces of neurites, forming a barrier between cells even in the absence of an ultrastructurally visible basement membrane. In the context of fasciculation, this barrier might prevent neurites from contacting inappropriate partners to which they would promiscuously adhere. In support of this view, we note that, in the case of IL2 defasciculation and axon flip-over, the neurites do not appear to lose adhesivity but instead they become tightly adhered to incorrect partners. In the context of preventing cellular breakage, this barrier would protect neurons and glia from neighboring structures that would otherwise impinge on them and cause fragmentation or rupture. In support of this view, we note that the BAG dendrite appears to break at a stereotyped position where it is normally thinner, as if under pressure from neighboring structures. It is less clear how this would cause fragmentation of the amphid sheath glial cell, but possibilities include aberrant adhesion with inappropriate neighbors or leakage through a perforated ECM that could cause shedding of small pieces of the glial cell as the animal grows and moves.

Consistent with the idea that DIG-1 contributes to basal ECM barriers between neurons, ultrastructural analysis has revealed defects in the basement membrane of *dig-1* mutants [12]. By contrast, DYF-7 is a component of apical ECM that lines the lumen of developing epithelial tubes [9]. This suggests a surprising interplay between basal and apical ECM during IL2 dendrite extension. One speculative possibility is that, while apical ECM acts in the lumen of developing epithelial (or glial) tubes to prevent rupture [35], basal ECM creates a protective barrier around the structure that reduces the overall mechanical stress it experiences during morphogenesis.

A perplexing feature of *dig-1* phenotypes is their diversity and cell-type specificity – some ventral cord axons are affected more than others [12], BAG undergoes breakage while a similar neuron, URX, is unaffected (our unpublished observations), and IL2 neurons are affected during embryonic development while other cell types are only affected near adulthood [11,12]. One possibility is that these phenotypic differences reflect the local environment around each cell, including the forces exerted on it throughout development by neighboring structures. Overall, this model suggests that DIG-1 may play a role in preventing inappropriate cell-cell interactions that is as important to nervous system organization as promoting the correct contacts.

## METHODS

### Strains and strain maintenance

All strains were constructed in the N2 background and cultured at 20°C on nematode growth medium (NGM) plates seeded with *E. coli* OP50. Strains, transgenes, and alleles used in this study are listed in Supplementary Tables S1-S3.

### Forward genetic screens and genetic mapping

All mutants were isolated from F2 nonclonal forward genetic screens. Briefly, L4 animals carrying an integrated cell-specific reporter (amphid sheath: *F16F9.3*pro:mCherry; BAG: *flp-17*pro:GFP; IL2: *klp-6*pro:GFP) were mutagenized for 4 h at room temperature in 70 mM ethyl methansulfonate (EMS, Sigma). Nonclonal F2 progeny were visually inspected for altered cellular morphologies using a Nikon SMZ1500 stereomicroscope with an HR Plan Apo 1.6× objective. Animals with aberrant cell shapes were recovered to individual plates. *hmn152, hmn158, hmn227,* and *hmn259* mutations were crossed to the Hawaiian wild strain CB4856 and mapped to the *dig-1* locus by whole-genome sequencing of pooled recombinants [36,37]. The sequence mutations identified in each allele are listed in Supplementary Table 2.

### Microscopy and image processing

Animals were immobilized with 10-100 mM sodium azide dissolved in M9, depending on developmental stage, mounted on a 2% agarose pad, and covered with a No. 1.5 coverslip. Spherical aberration was minimized using immersion oil matching. Z-stacks were acquired using a DeltaVision Core deconvolution imaging system (Applied Precision) with the InsightSSI light source; UApo 40x/1.35 NA oil immersion objective, PlanApo 60x/1.42 NA oil immersion objective, or UPlanSApo 100x/1.40 NA oil immersion objective (Olympus); the standard DeltaVision live cell excitation and emission filter set; and a Photometrics CoolSnap HQ2 CCD camera (Roper Scientific). Images were acquired and deconvolved with Softworx 5.5 (Applied Precision). Images are displayed as maximum intensity projections generated in Priism [38]. Images were pseudocolored and image brightness was linearly adjusted using Adobe Photoshop. IL2 dendrite lengths were measured using the Segmented Line tool in Fiji (NIH). To control for differences in head size, dendrite lengths are normalized to the distance from the cell body to the nose tip. P-values were generated using the Wilcoxon Rank-Sum (Mann-Whitney U) test in RStudio.

## ACKNOWLEDGEMENTS

We gratefully acknowledge WormBase; the CGC, which is funded by NIH Office of Research Infrastructure Programs (P40 OD010440); and funding from the NSF Graduation Research Fellowship Program (E.R.C), National Institutes of Health (F31NS103371 to E.R.C; R01GM108754 and R01NS112343 to M.G.H.), and the Hearst Fellows Program at Harvard University (K.M.).

## Notes

### Competing Interest Statement

The authors have declared no competing interest.

