## Supplemental Material for "Loss of the extracellular matrix protein DIG-1 causes glial fragmentation, dendrite breakage, and dendrite extension defects"

**Table S1. Strains used in this study**

| Strain | Genotype | Figure(s) |
|---|---|---|
| CHB1652 | *dig-1(hmn227)* III*; hmnIs13* | 1 |
| CHB1166 | *dig-1(hmn152)* III*; oyIs82* X | 1 |
| CHB2250 | *dig-1(hmn158)* III*; oyIs82* X | 1 |
| CHB3008 | *dig-1(hmn259) myIs13* III | 1 |
| CHB1549 | *hmnIs13* | 2 |
| CHB4342 | *dig-1(n1321)* III*; hmnIs13* | 2 |
| PY8503 | *oyIs82* X | 3 |
| CHB4343 | *dig-1(n1321)* III*; oyIs82* X | 3 |
| PT2762 | *myIs14* IV | 4, 5 |
| CHB2940 | *dig-1(n1321)* III*; myIs14* IV | 4, 5 |
| CHB2937 | *myIs14* IV; *dyf-7(ns119)* X | 5 |
| CHB2944 | *dig-1(n1321)* III*; myIs14* IV; *dyf-7(ns119)* X | 5 |

**Table S2. Transgenes used in this study**

| Transgene | Description | Reference |
|---|---|---|
| *hmnIs13* | *F16F9.3*pro:mCherry, *grl-2*pro:YFP, *gcy-8*pro:CFP | Mizeracka et al., bioRxiv 2019 |
| *oyIs82* | *flp-17*pro:GFP, *unc-122*pro:dsRed | Gift of Astrid Cornils and Piali Sengupta |
| *myIs13* | *klp-6*pro:GFP | Schroeder et al., Curr. Biol. 2013 |
| *myIs14* | *klp-6*pro:GFP | Schroeder et al., Curr. Biol. 2013 |

**Table S3. Alleles generated in this study**

| Allele | Sequence |
|---|---|
| *dig-1(hmn152)* | CTGATGGT[C>T]AGATTCTT |
| *dig-1(hmn158)* | ATGGACAG[g>a]tacgatta |
| *dig-1(hmn227)* | ATGGACAG[g>a]tacgatta |
| *dig-1(hmn259)* | GGTTCAC[C>T]CAATAACA |

Substitutions are bracketed. Uppercase corresponds to predicted exons; lowercase to predicted introns. *hmn158* and *hmn227* were isolated from independent screens but cause an identical nucleotide change.
